# Ecosystem engineering alters density-dependent feedbacks in an aquatic insect population

**DOI:** 10.1101/2021.01.12.426351

**Authors:** Joseph S. Phillips, Amanda R. McCormick, Jamieson C. Botsch, Anthony R. Ives

## Abstract

Ecosystem engineers have large impacts on the communities in which they live, and these impacts may feed back to populations of engineers themselves. In this study, we assessed the effect of ecosystem engineering on density-dependent feedbacks for midges in Lake Mývatn, Iceland. The midge larvae reside in the sediment and build silk tubes that provide a substrate for algal growth, thereby elevating benthic primary production. Benthic algae are in turn the primary food source for the midge larvae, setting the stage for the effects of engineering to feed back to the midges themselves. Using a field mesocosm experiment manipulating larval midge densities, we found a generally positive but nonlinear relationship between density and benthic production. Furthermore, adult emergence increased with the primary production per midge larva. By combining these two relationships in a simple model, we found that the positive effect of midges on benthic production weakened the negative density dependence at low to intermediate larval densities. However, this benefit disappeared at high densities when midge consumption of primary producers exceeded their positive effects on primary production through ecosystem engineering. Our results illustrate how ecosystem engineering can alter density-dependent feedbacks for engineer populations.

## Introduction

Ecosystem engineering is a class of ecological interactions whereby one population affects others through alterations to the physical environment (Jones et al., 1994; Wilby, 2002). Like all interspecific interactions, ecosystem engineering has the potential to generate feedbacks among various members of a community (Bertness and Leonard, 1997; Largaespada et al., 2012; Donadi et al., 2014; Sanders et al., 2014). For example, physical structure provided by coral can ameliorate competition with algae by benefiting grazers that reduce algal abundance (Bozec et al., 2013). As ecosystem engineers are (by definition) the source of engineering effects within an ecosystem, feedbacks between engineering and the engineers themselves are central to the dynamical consequences of engineering for the community as a whole (Hastings et al., 2007; Sanders et al., 2014).

To understand the role of engineering feedbacks for the population dynamics of ecosystem engineers, it is useful to relate those feedbacks to the strength of density dependence (Hastings et al., 2007; Cuddington et al., 2009). Using a simple mathematical model, Cuddington et al. (2009) showed that a wide range of dynamical behavior is possible for populations of ecosystem engineers, including stable persistence, extinction, unbounded growth, and alternative states. The dependence of engineering effects on population density and the subsequent feedback of engineering to density dependence are key factors determining the overall dynamical consequences of ecosystem engineering. Despite their theoretical importance, quantitative characterizations of density-dependent engineering feedbacks for natural populations are limited. While previous studies have established the existence of engineering-mediated feedbacks to engineer populations (e.g., Largaespada et al., 2012; Bozec et al., 2013; Donadi et al., 2014), they have generally not done so across a range of engineer densities as is required to directly quantify the density dependence of population growth.

We assessed the effect of ecosystem engineering on the sign and magnitude of density-dependent survival and emergence of the midge *Tanytarsus gracilentus* (Diptera: Chironomidae) in Lake Mývatn, Iceland. The larvae of *T. gracilentus* dwell in the sediment and build silk tubes that elevate primary production by providing a substrate for algal growth (Herren et al., 2017; Phillips et al., 2019), similar to other aquatic macroinvertebrates (Largaespada et al., 2012; Donadi et al., 2014; Hoelker et al., 2015). The larvae feed on benthic algae (mainly diatoms), which means that their enhancement of benthic production may benefit their own survival, emergence, and subsequent reproduction (Ingvason et al., 2004). However, midge consumption may also reduce algal biomass, potentially leading to intraspecific competition and negative density dependence (Einarsson et al., 2016). Indeed, the *T. gracilentus* population in Mývatn shows large fluctuations in abundance that are likely driven by food limitation, although these fluctuations cannot be explained purely in terms of classical consumer-resource cycles (Ives et al., 2008). Characterizing the nature of density dependence is important for understanding the complex population dynamics of *T. gracilentus*, making it a valuable case for exploring the effects of ecosystem engineering on density dependence of engineer populations.

To evaluate the role of ecosystem engineering on density-dependent survival and emergence in *T. gracilentus*, we conducted a field mesocosm experiment across a range of experimental larval densities. This allowed us to directly quantify (i) the relationship between benthic primary production and larval midge density and (ii) the relationship between adult emergence rates and primary production per larval midge. We then combined these two relationships with a simple model that allowed us to isolate the contribution of larval midge effects on primary production to their density-dependent survival and emergence.

## Methods

Mývatn is a large (37m^2^), shallow (mean depth: 2.5m), naturally eutrophic lake in northeastern Iceland (65°40’N 17°00’W) (Einarsson et al., 2004). It is separated into two ecologically distinct basins (north and south). Our study was conducted in 2017 at three south basin sites (E2, E3, and E5) with soft substrate that were selected to represent a range of ecological conditions and *T. gracilentus* abundance (Figure S1). In sediment cores taken throughout the summer of 2017, E3 had the highest Tanytarsini (including *T. gracilentus*) densities (mean ± standard error: 69, 058 ± 14, 595 m^−2^), followed by E5 (30, 648 ± 11, 767), and then E2 (431 ± 172). Maximum densities in Mývatn have exceeded 500, 000 m^−2^ (Thorbergsdóttir et al., 2004). In the summer of 2017, E2 was subject to an expanding mat of filamentous green algae (Cladophorales) that was largely absent from E3 and E5. Furthermore, E2 was consistently colder (mean difference 1°C) than E3 and E5 during the experiment period (Figure S1). In contrast, photosynthetically active radiation (PAR) was similar among the sites, due to their similar depths (E2: 2.8m; E3: 3.3m; E5: 2.6m) and water clarities throughout the south basin in 2017. Light and temperature data were collected with two loggers (HOBO Pendant, Onset Computer Corporation) deployed on the lake bottom at each site and set to log every 30 minutes. Light was measured as visual intensity and approximately converted to PAR using a standard correction (Thimijan and Heins, 1983).

We conducted our field mesocosm experiment using a design similar to Phillips et al. (2019). On 28 June 2017, we collected sediment cores from the three study sites using a Kajak corer. For each site, we pooled the sediment from the different cores while keeping the top 5cm (“top”) and next 10cm (“bottom”) separate. We then sieved the sediment through either 125 (top) or 500*μ*m (bottom) mesh to remove midge larva; sieving also removed the surface Cladophorales abundant in cores from E2. The sediment was left to settle for 4d in a cool and dark location. We constructed the mesocosms by stocking the sediment into clear acrylic tubes (33cm height × 5cm diameter) sealed from the bottom with foam stoppers. We first added 10cm of bottom sediment and then 5cm of top sediment, to mimic the layering in the lake. The sediment layer of each mesocosm was wrapped with 4 layers of black plastic to eliminate light from the sides.

On 3 July, we took sediment cores at E3 and sieved them through 125*μ*m mesh to collect Tanytarsini larvae; the vast majority were likely *T. gracilentus*, although identification to the species level could not readily be done on live individuals. Tanytarsini progress through four instars before emerging as adults. We attempted to select individuals the general size of second instar larvae to maximize the duration of the experiment before emergence. On 4 July, we stocked the mesocosms with four densities of Tanytarsini larvae: 0, 50, 100, 200 per mesocosm (0, 25000, 51000, and 102000 m^−2^). Each site × density combination had four replicates, for a total of 48 mesocosms. We filled the mesocosms with water collected from near Mývatn’s southern shore and gave the midges 24h to settle before deploying the mesocosms in the lake. On 5 July, we distributed the mesocosms corresponding to each site onto two racks and then deployed them at their respective sites on the lake bottom. The tops of the mesocosms were left open to allow exchange between the mesocosms and the lake water column.

On 10 and 11 July (days 5 and 6), we estimated gross primary production (GPP) in the mesocosms by measuring the change in dissolved oxygen (DO) concentration during sealed incubations (similar to Phillips et al., 2019). The incubations were conducted in situ at the respective sites to incorporate spatial variation in ambient conditions, such as light and temperature. Each mesocosm was first incubated under ambient light to give an estimate of net ecosystem production (NEP), followed by a dark incubation with the top of each mesocosm wrapped in 4 layers of black plastic to give an estimate of ecosystem respiration (ER). NEP + ER gives an estimate of GPP, assuming that ER is the same during both the light and dark incubations. Half of the mesocosms at each site were incubated on 10 July, while the other half were incubated on 11 July; all of the mesocosms incubated on a given day for a given site were on the same experimental rack and so constituted a block. The incubations lasted 3–5h, and the tops of the mesocosms were sealed with rubber stoppers for the duration. DO was measured using a handheld probe (ProODO, YSI, Yellow Springs, Ohio, USA), and we gently stirred the water within each mesocosm to homogenize it before taking the reading. We repeated the incubation procedure on 21 and 23 July (days 16 and 18). Due to difficult weather, we were unable to perform the incubations at the respective sites. Therefore, on 21 July all of the mesocosms were moved to a bay on the southern shore of the south basin (depth ≈ 1.7m). The light incubations lasted 3–5h, while the dark incubations lasted 4–10h. While variation in incubation duration was not ideal, the DO in the dark incubations remained above anoxic conditions (minimum DO >10 mg L^−1^). We converted GPP to units of mg O_2_ m^−2^ h^−1^, accounting for incubation duration and water column depth within each mesocosm.

On 23 July, shortly before the expected time of midge emergence, we removed the mesocosms from the lake and covered the top of each with mesh to catch adult midges as they emerged. We kept the mesocosms outdoors in baths of cold tap water to moderate temperature. Every 1–3d for the next 13d, we collected the emerging adults from the mesocosms. While these were not individually identified, the vast majority appeared to be Tanytarsini. Furthermore, there was a strong association between the number of Tanytarsini larvae stocked in the mesocosms and the number of adults that emerged (Spearman rank correlation of 0.82; *P* < 0.0001).

We quantified the relationship between GPP and larval density using a linear mixed model (LMM). The model included initial density (numeric), site (three levels), incubation day (two levels; either days 5–6 or 16–18), and their two-way interactions as fixed effects. Because we expected the relationship between GPP and initial density to be nonlinear, we also included 2nd and 3rd order polynomial terms for initial density (without any interactions) in the model. We chose a third degree polynomial because this gave the same number of parameters to estimate as would have been the case if each of the four density levels were treated categorically (including the intercept). The polynomial regression allowed us to treat density as a numeric variable and simplify the model by only including interactions with the linear density term. We accounted for variation in ambient conditions during the incubations by including linear terms for PAR and temperature estimated for each block at each site. Finally, we included random effects for experimental rack and mesocosm identity to account for blocking and repeated measures.

We used a binomial generalized linear mixed model (GLMM) to analyze variation in the number of midges that emerged as adults relative to the initial number of larvae (excluding the zero treatment). We included initial density (numeric), site (three levels), and their interaction as fixed effects. We included random effects for experimental rack and mesocosm identity to account for blocking and potential overdispersion, respectively; the latter was equivalent to assuming the residuals followed a logit-normal binomial distribution.

Our goal in this study was to assess the effect of midge ecosystem engineering on their emergence. To do this, we fit a logit-normal binomial GLMM to the number of midges that emerged as adults relative to the initial number of larvae with GPP per initial larva as the sole fixed effect. We then projected the number of emerging adult under two scenarios: (i) using predicted values of GPP as a function of site, incubation period, and larval density treatment according to the polynomial LMM described above, and (ii) using predicted values of GPP as a function of site and incubation period, but with larval density set to zero (on the natural scale). Scenario (ii) implies that GPP per larva declines across the midge treatments due purely to the greater number of larvae over which production is distributed. Scenario (i) differs from scenario (ii) by also including direct effects of larval density on GPP. The difference between scenarios (i) and (ii) gives a measure of the effect of larval density on the density dependence of survival and emergence with and without the effects of midge larvae on GPP.

Statistical analyses were conducted in R 4.0.3 (R Core Team, 2020), using the lme4 package to fit the LMM and GLMMs (Bates et al., 2015). We calculated *P*-values with *F*-tests using the Kenward-Roger correction for the LMM and parametric-bootstrapped likelihood-ratio tests (LRTs) based on 2000 simulations for the GLMMs (`simulate` function in the native `stats` package). We report both Type III and Type II tests unless otherwise noted, to balance concerns of inflated Type I errors that can occur when dropping terms with the poor statistical inference that can come from overparameterized models.

## Results

The mesocosm experiment revealed a nonlinear relationship between GPP and larval density, as the first, second, and third-degree terms associated with midge larval density were all statistically significant (Table I). This relationship was generally positive, although it saturated and was possibly negative at the highest densities (Figure 1). On days 5–6 of the experiment, all three sites had similar GPP-density relationships and overall levels of GPP. However, there were statistically significant day × site, day × density, and site × density interactions that manifested as differences between the sites on days 16–18. For sites E2 and E5, the GPP-density relationship was weaker on days 16–18 than on days 5–6, while the density effect at E3 remained largely similar through time. Furthermore, GPP for E3 and E5 was higher on days 16–18 than on days 5–6, while for E2 it was lower. These relationships corrected for the positive effect of ambient temperature during the measurement incubations. Therefore, the temporal patterns likely reflect real divergences between the productivity of the mesocosms at the three sites through time, rather than transient differences in ambient environmental conditions during the measurements.

**Table I:**
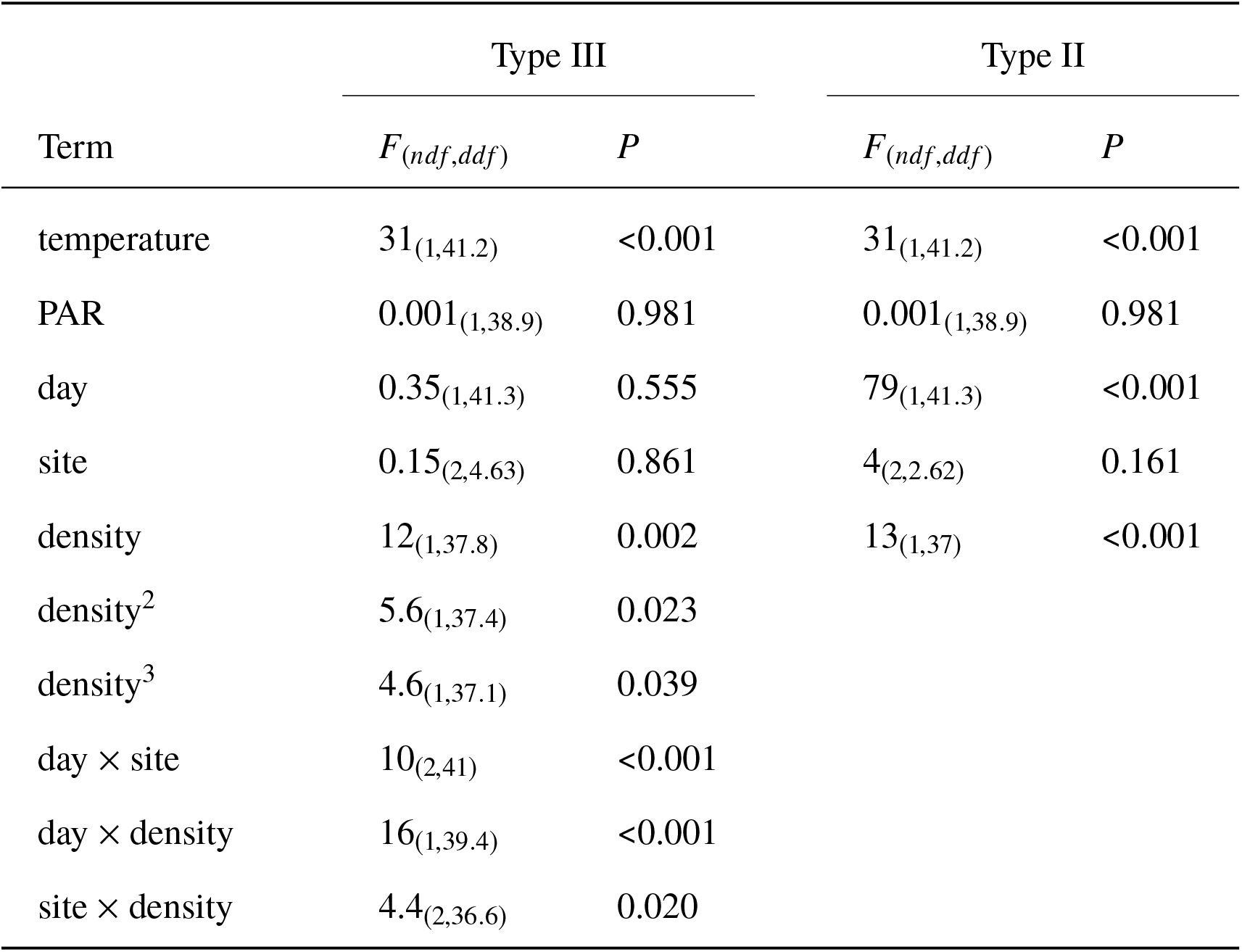
LMM for mesocosm GPP. *P*-values are from *F*-tests with the Kenward-Roger correction. The model included random effects for block (σ = 0.001) and mesocosm identity (σ = 0.013), with residual standard deviation σ = 0.012. The factor “day” had two levels (either day 5–6 or 16–18).

**Figure 1.**
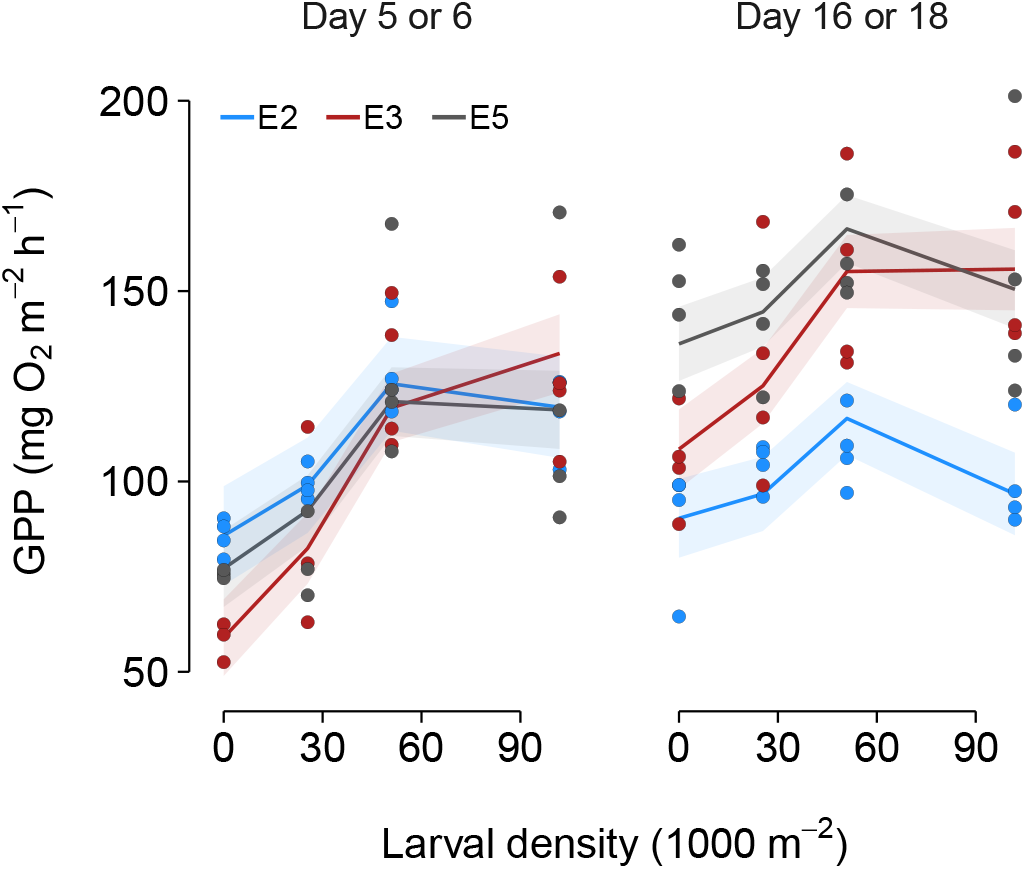
GPP as a function of initial larval density. The points show the observed data standardized to the mean water temperature measured during the incubations. The solid lines show fitted values from the third-degree polynomial LMM. While the model provides a smooth fit to the data, we connected the fitted values at the experimental densities with straight lines to draw attention to discrete levels at which the measurements were taken. The shaded regions show the standard errors estimated from the covariance matrix associated with the model fit.

The proportion of individuals that emerged as adults declined with initial larval density (Type II LRT: 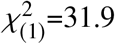; *P* < 0.001), indicating negative density dependence. Neither the main effect of site (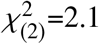; *P* = 0.422) nor its interaction with density (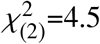; *P* = 0.147) were statistically significant. Proportional emergence increased with the GPP per initial larva (LRT: 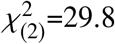; *P* < 0.001; Figure 2), which is consistent with the hypothesis that negative density dependence is related in part to food limitation.

**Figure 2.**
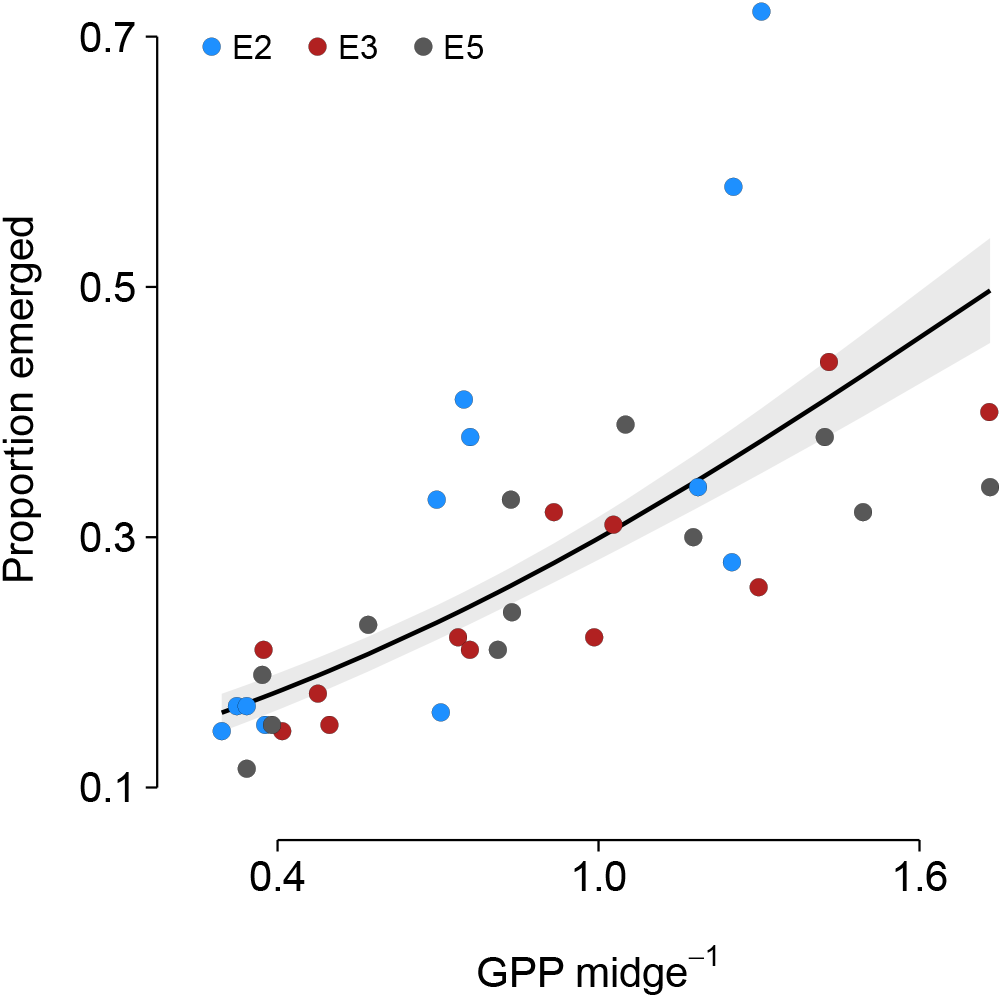
Adult emergence rate as a function of GPP (mg O_2_ m^−2^ h^−1^) per initial midge larva. The points show the observed data, the line shows the fitted values from a GLMM, and the shaded region shows the standard errors estimated from the covariance matrix associated with the model fit.

We assessed the effect of larval density on adult emergence by projecting the number of emerging midges under two scenarios: (i) including the effect of larval density on overall GPP and (ii) assuming that GPP was constant and therefore was not affected by midge larvae. Differences between the two scenarios quantified the consequences of midge effects on GPP for density-dependent emergence. The positive effect of larvae on GPP reduced the negative effect of larval density on the proportion of individuals that emerged as adults (Figure 3a). However, because the midge effect on GPP plateaued at high initial midge density (Figure 1), proportional emergence at high density converged on what it would be without the midge effect on GPP. There was modest variation in the midge effect on density dependence among the three sites, with the effect being greatest at E3. This reflects the fact that the positive midge effect on GPP declined through time at E2 and E5, while at E3 it remained relatively constant. The nonlinear effect of midge larvae on GPP resulted in a hump-shaped relationship between total emergence and larval density (Figure 3b), indicating that the greatest emergence occurred at intermediate densities.

**Figure 3.**
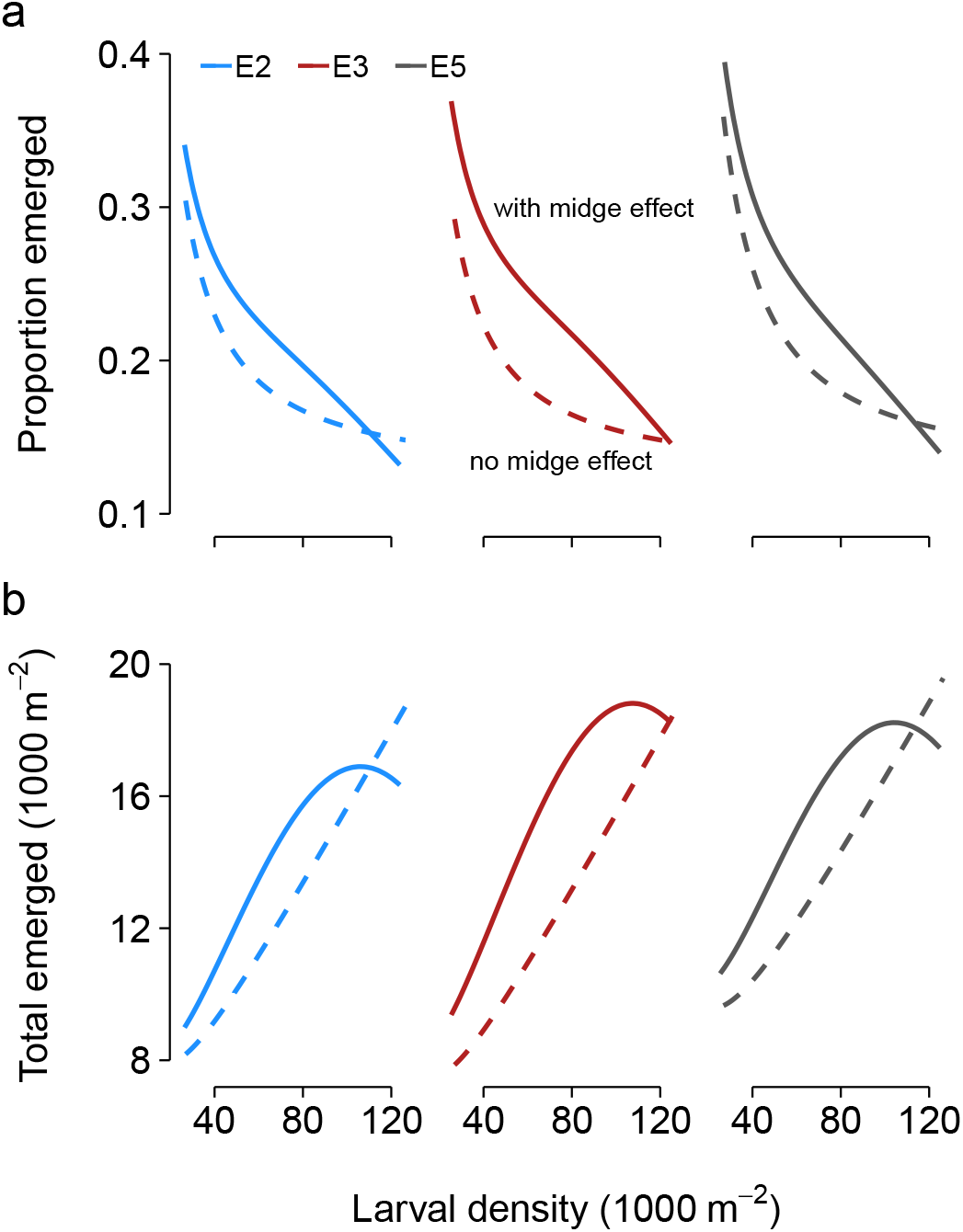
Consequences of midge effects on GPP for negative density dependence of either (a) the proportion of larvae that emerged as adults or (b) the total number that emerged. The curves are derived from the conjunction of the LMM in Figure 1 and the GLMM in Figure 2. The solid lines (“with midge effect”) show modeled emergence rates including the modeled effect of midges on overall GPP, while the dashed lines (“no midge effect”) exclude midge effects on overall GPP. The difference between the two lines is a measure of the midge effect on density-dependent emergence as mediated by midge effects on GPP.

## Discussion

Our field mesocosm experiment showed that an ecosystem engineer population has positive effects on its food resources, thereby establishing a positive feedback that alters density-dependent survival and emergence. We found that Tanytarsini larvae in Mývatn have large positive effects on benthic primary production, which previous studies have shown are driven at least in part by physical structure provided by the silk tubes in which the larvae reside (Hoelker et al., 2015; Phillips et al., 2019). Because midge larvae feed on benthic diatoms (Ingvason et al., 2004), stimulation of benthic production increased the amount of food per individual midge, which in turn weakened negative density-dependence arising from food limitation at moderate midge densities. However, the amelioration of negative density dependence largely disappeared at high densities, presumably due to midge suppression of diatom growth through consumption.

The nonlinear feedback of midge engineering on primary production meant that the weakening of density dependence in emergence was greatest at intermediate densities. Given the suppression of algal biomass through grazing (Einarsson et al., 2016), it is likely that the effect of midges on production becomes negative at the highest densities observed in the lake. This suggests that while the feedback through midge engineering weakens negative density dependence at moderate densities, this is not enough to overcome the consumptive effect of midges at high densities. The nonlinearity of ecosystem engineer effects has previously been identified as an important factor in governing the effects of engineering on community dynamics (Bozec et al., 2013). Despite weakening density dependence at low to moderate densities, midge engineering did not lead to positive density dependence in per capita emergence. However, the nonlinear engineering feedback did result in a hump-shaped relationship between total emergence and larval density, which could lead to overcompensatory dynamics (Turchin, 2003; Cuddington et al., 2009). It is important to note that our study does not include information on reproduction and so does not fully capture the effect of density dependence on the full life cycle. Nonetheless, given the short duration of the adult stage and high egg production per adult *T. gracilentus*, it is possible that the dynamics of the “engineering stage” (i.e., larvae) are principally driven by larval survival, of which adult emergence is a direct extension. While ecosystem-engineering effects of other benthic invertebrates may differ from tube-building midges by being concentrated in the adult stage (e.g. mussels; Largaespada et al., 2012), juvenile production may be similarly unlimited by adult abundance such that survival of the engineering stage is most relevant for their dynamics.

The midge effect on benthic productivity across the three sites was similar at the beginning of the experiment, but diverged through time even after accounting for variation in ambient conditions during the productivity measurements. Furthermore, the engineering feedback subtly differed between sites, with midge emergence experiencing the greatest benefit from engineering at E3, which was the site with the greatest ambient densities at the beginning of the experiment. This suggests that local environmental context altered the consequences of ecosystem engineering, and these alterations had the potential to persist through time. While our experiment was not directly able to test for such legacies, plausible candidate causes are temperature (which was consistently lower at E2), and sediment nutrient concentrations which vary across locations. These environmental variables could have altered algal community composition or abundance (Gudmundsdottir et al., 2011; McCormick et al., 2019), thereby mediating their capacity to respond to the amelioration of light limitation provided by midge engineering (Phillips et al., 2019). Various studies have identified the role of environmental variation in mediating the strength and sign of ecosystem engineering (Wright et al., 2006; Lathlean and McQuaid, 2017). When the effects of environmental mediation persist through time, they may decouple the dynamics of the engineers and their community-wide effects. Such decoupling fundamentally alters the density-dependence of engineering feedbacks, and therefore may play a central role in determining the dynamics of ecosystem engineering (Cuddington et al., 2009).

## Acknowledgments

This work was supported by National Science Foundation grants DEB-1052160, DEB-1556208 to Anthony R. Ives, and Graduate Research Fellowships DGE-1256259 and DGE-1747503. The Mývatn Research Station directed by Árni Einarsson provided logistical and scientific support. We thank Kristian Riley Book, Natalie Schmer, Bethany Smith, and Aspen Ward for assistance with fieldwork.

---

**Figure S1:**
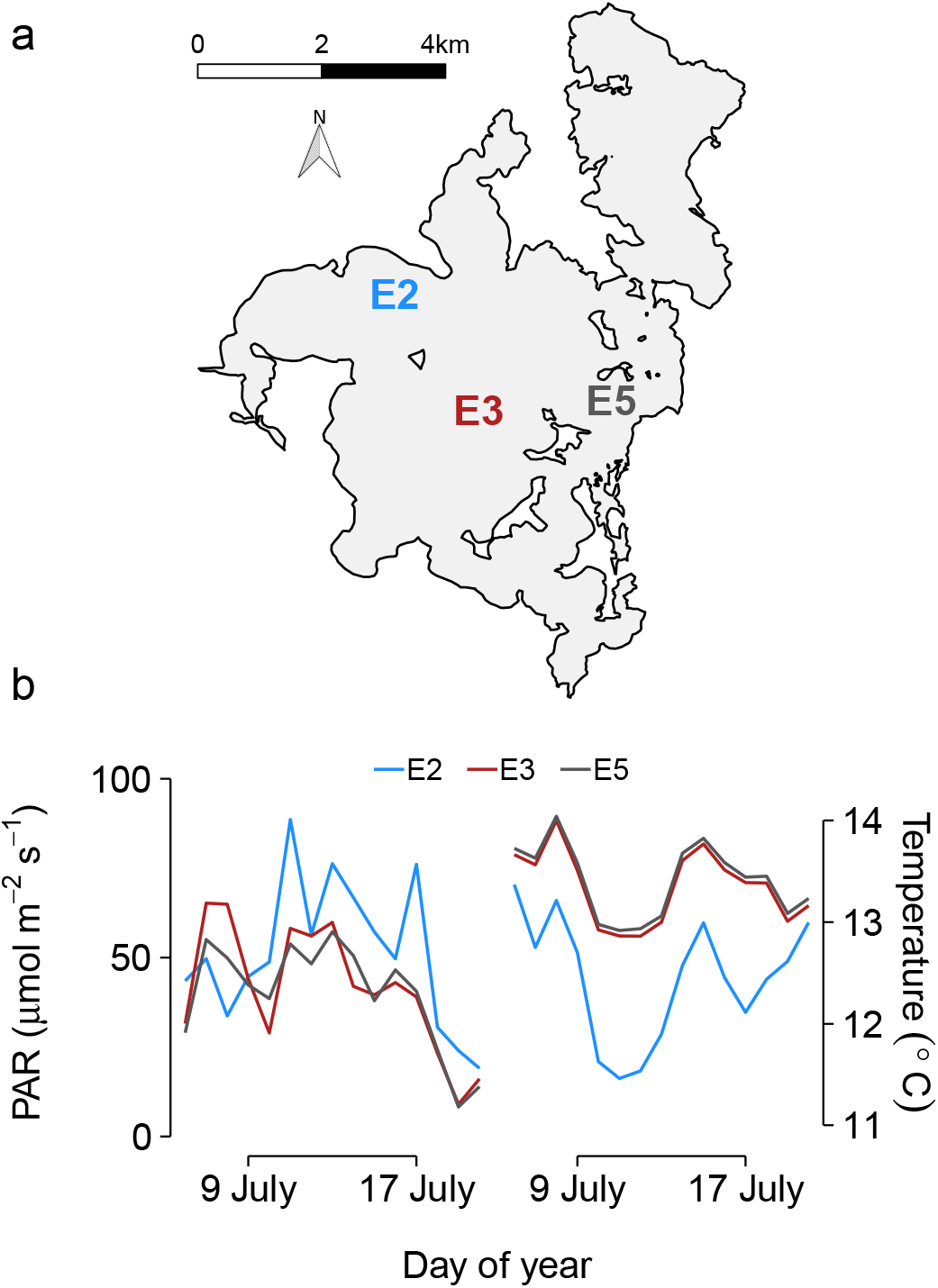
Experimental sites. (a) The three sites covered a wide range of Mývatn’s south basin. Light gray areas indicate water, while white areas indicate land. (b) Mean daily PAR and temperature were calculated by averaging half-hourly measurements from two loggers deployed on the lake bottom for each site.

